# Sentinel plants enable quantitative monitoring of bioavailable nitrate in soils and microbial environments

**DOI:** 10.64898/2026.03.18.712767

**Authors:** Eugene Li, Chiara Berruto, Tufan Oz, Elisa Grillo, Catherine Griffin, Yunqing Wang, Jolie W. Jones, Gozde S. Demirer

**Affiliations:** Chemistry and Chemical Engineering Division, California Institute of Technology (Caltech), Pasadena, California, USA 91125; Biology and Biological Engineering Division, California Institute of Technology (Caltech), Pasadena, California, USA 91125

**Keywords:** plant biosensors, sentinel plants, nitrate, nitrogen cycling, synthetic biology, rhizosphere, plant-microbe interactions

## Abstract

Microbial transformations of nitrogen in soils strongly influence plant nutrition and ecosystem function, yet these processes remain difficult to monitor. Existing approaches rely largely on destructive soil sampling and laboratory analysis, limiting the ability to track nitrate dynamics *in situ.* Here, we engineer “sentinel plants,” genetically encoded plant biosensors that convert nitrate perception into a quantitative signal reporting plant-accessible nitrate. The sensor uses a synthetic nitrate-responsive promoter coupled to a ratiometric luciferase reporter, enabling high-dynamic-range measurements. Sentinel plants exhibit a dose-dependent, reversible nitrate response with high specificity over alternative nitrogen sources. In agricultural soils from multiple California field sites, sensor output closely tracked analytically measured nitrate concentrations and resolved nitrate amendments without destructive extraction. Beyond environmental sensing, sentinel plants enabled screening of nitrogen-fixing microbial communities and the detection of microbially generated nitrate in both liquid culture and soil systems. Using this platform, we identified a minimal three-member microbial consortium capable of converting atmospheric nitrogen into nitrate via sequential nitrogen fixation and nitrification. This consortium increased tissue nitrate accumulation and plant fresh weight, demonstrating that sentinel plants can both monitor nitrate availability and identify microbial communities that enhance plant growth.

**Significance Statement:** Nitrogen availability in soils fluctuates across space and time, yet most measurements rely on extracting soil samples and analyzing them in the laboratory. Such measurements provide only snapshots of nitrogen status and do not necessarily reflect the nitrogen that plants themselves experience. Here, we engineer plants that act as living nitrate sensors by converting nitrate perception into a measurable optical signal. Because these sensors operate within intact plants, they report nitrate availability as integrated through plant uptake and physiology rather than through chemical extraction alone. Using this platform, we tracked nitrate levels in agricultural soils and identified a minimal microbial consortium capable of converting atmospheric nitrogen into plant-available nitrate. This plant-based sensing strategy enables direct monitoring of nitrogen dynamics in soils and microbial environments, providing a platform for identifying microbial communities that enhance nitrogen availability for crops.

## Introduction

Nitrogen availability strongly limits plant productivity and global agricultural output, with nitrate representing the dominant bioavailable nitrogen source in many aerobic soils. Optimizing nitrate availability is therefore central to both crop yield and environmental stewardship. Excess nitrogen fertilization promotes nitrate leaching, eutrophication, and nitrous oxide emissions, whereas insufficient nitrogen reduces crop productivity^1^. These competing pressures necessitate precise sensing strategies that quantify bioavailable nitrate with sufficient spatial and temporal resolution to guide nitrogen management decisions.

Most nitrate measurements rely on destructive extraction-based assays in which soil samples are collected, chemically extracted, and quantified using laboratory techniques, such as cadmium-reduction colorimetry^2^ or ion chromatography^3^. Although accurate, these approaches require laboratory infrastructure and capture only a snapshot of nitrate levels at a specific time and location. In agricultural soils, nitrate exhibits strong spatial and temporal heterogeneity driven by processes such as irrigation, nitrification, denitrification, and plant uptake. As a result, sparse sampling often fails to capture nitrate dynamics that are agronomically and environmentally important. Field test kits and portable colorimetric assays reduce cost and complexity but still require physical sampling and extraction^4–6^, and therefore provide discrete measurements rather than continuous or spatially resolved monitoring.

Electrochemical sensing approaches, particularly nitrate ion-selective electrodes, offer an alternative strategy for in-field measurements^7–11^. These systems enable rapid on-site sensing and can provide higher spatial and temporal sampling density than laboratory workflows. However, electrochemical sensing in soils remains challenged by signal drift, cross-sensitivity to interfering ions, and instability of reference electrodes in chemically and physically heterogeneous soil matrices. Fluctuations in temperature and ionic strength can further compromise calibration stability, often requiring frequent recalibration and limiting long-term autonomous monitoring. Moreover, remote sensing using unmanned aerial vehicles equipped with multispectral or hyperspectral sensors is increasingly used to support nitrogen management strategies^12–14^. These systems typically estimate nitrogen status indirectly through canopy spectral responses, such as chlorophyll-related reflectance indices. Although hyperspectral imaging has enabled advances in predictive modeling, challenges remain, including limited model transferability across cultivars, growth stages, and environmental conditions. Methods capable of directly quantifying plant-accessible nitrate in soil environments, therefore, remain an important unmet need.

Biological sensing strategies offer a complementary paradigm in which living systems transduce environmental nitrate exposure into measurable outputs^15^. Whole-cell microbial biosensors have been developed to report nitrate availability in environmental matrices. For example, bacterial reporters have been engineered in which nitrate-responsive promoters controlled the expression of optical reporters, enabling detection of nitrate gradients in soil and rhizosphere environments^16^. However, microbial biosensors report the exposure of microbes to nitrate rather than the availability of nitrate to plants. Because microbes and plant roots occupy distinct spatial niches and differ in nutrient uptake dynamics, nitrate concentrations sensed by microbes may diverge from those accessible to roots. In addition, deploying engineered microbial reporters requires maintaining living strains in complex soil communities, which can complicate reproducibility, interpretation, and regulatory deployment.

Plants themselves directly integrate soil nitrate through root uptake and internal transport, linking environmental nitrogen availability to whole-plant physiology. Consequently, signals measured in plant tissues could provide a plant-relevant readout of soil nitrate dynamics without destructive soil sampling. Early work established a synthetic nitrate-responsive promoter (NRP) that enabled nitrate-responsive transcriptional reporters and provided a foundational platform for developing plant-based nitrate sensors^17^. However, the sensor output was attenuated by ammonium, limiting specificity for nitrate in complex environments. Several additional genetically encoded nitrate sensors have been developed in plants to study nitrate transport and signaling within plant tissues. The fluorescent reporter NiTrac, derived from the nitrate transceptor NRT1.1, reports nitrate-dependent conformational changes associated with transporter activity^18^. Other protein-based sensors, such as NitraMeter3.0, derived from the bacterial nitrate-binding protein NasR, enable imaging of intracellular nitrate dynamics in plant cells^19^. More recently, the transcription factor NLP7 has been shown to function as a direct nitrate sensor that regulates primary nitrate signaling pathways in plants^20,21^.

These tools have provided important insights into plant nitrate signaling and intracellular nitrate dynamics and have enabled detailed mechanistic studies of nitrate perception and transport in plants. However, they were primarily designed for fluorescence-based measurements at cellular or tissue scales. Consequently, extending these sensing strategies to quantitative measurements of environmental nitrate availability across whole plants and heterogeneous soil environments remains challenging, and tools capable of reporting plant-accessible nitrate availability at the organismal level remain limited. This motivates the development of plant-integrated sensing platforms that convert endogenous nitrate perception into scalable whole-plant reporter outputs suitable for complex environments, complementing the existing cellular nitrate sensors. Such systems can enable direct observation of microbial nitrogen transformations that generate plant-accessible nitrate in soils, linking microbial activity in the rhizosphere to plant nutrient status.

Here, we engineer sentinel plants in which a synthetic NLP7-responsive promoter drives a ratiometric luciferase reporter, converting nitrate perception into a quantitative whole-plant signal. Because the reporter operates downstream of endogenous nitrate sensing, its output reflects the plant’s nitrate availability rather than concentrations measured in isolated soil extracts. Using this platform, we demonstrate quantitative sensing of nitrate across controlled media, agricultural soils, and microbial systems. As a proof of concept, we identified a three-member synthetic microbial consortium that converts atmospheric nitrogen into nitrate and enhances plant growth, with sentinel plant output quantitatively reporting nitrate generated by this community. These results illustrate how plant biosensors can enable the discovery and evaluation of microbial communities that influence plant nutrient acquisition, providing a generalizable framework for engineering plants as living sensors capable of monitoring plant-microbe-soil processes.

## Results

### Engineering and optimization of a genetically encoded nitrate sensor in Arabidopsis

Transcriptional interference between adjacent expression cassettes can compromise quantitative reporter measurements when inducible and normalizer reporters are encoded on the same plasmid^22,23^. To develop a ratiometric sensor capable of quantitatively reporting nitrate availability, we first optimized the cassette orientation. We used a previously characterized nitrate-responsive promoter, NRP^17^, to drive an enhanced luciferase (ELuc) reporter, together with a constitutive pNOS-driven RedF luciferase normalization cassette^24^.

The two transcriptional units were assembled in four relative orientations (**Fig. 1A**): head-to-head (H2H), head-to-tail (H2T), tail-to-head (T2H), and tail-to-tail (T2T), and tested in *Arabidopsis thaliana* mesophyll protoplasts exposed to nitrate starvation (0 mM), limited nitrate (1 mM), or sufficient nitrate (10 mM). Co-transfection (CoT) of separate ELuc and RedF plasmids was used as a control, representing minimal transcriptional interference. Among the configurations tested, the T2T orientation most closely reproduced the signal output and nitrate responsiveness observed in the CoT control (**Fig. 1B**) and was therefore used in all subsequent constructs.

**Figure 1.**
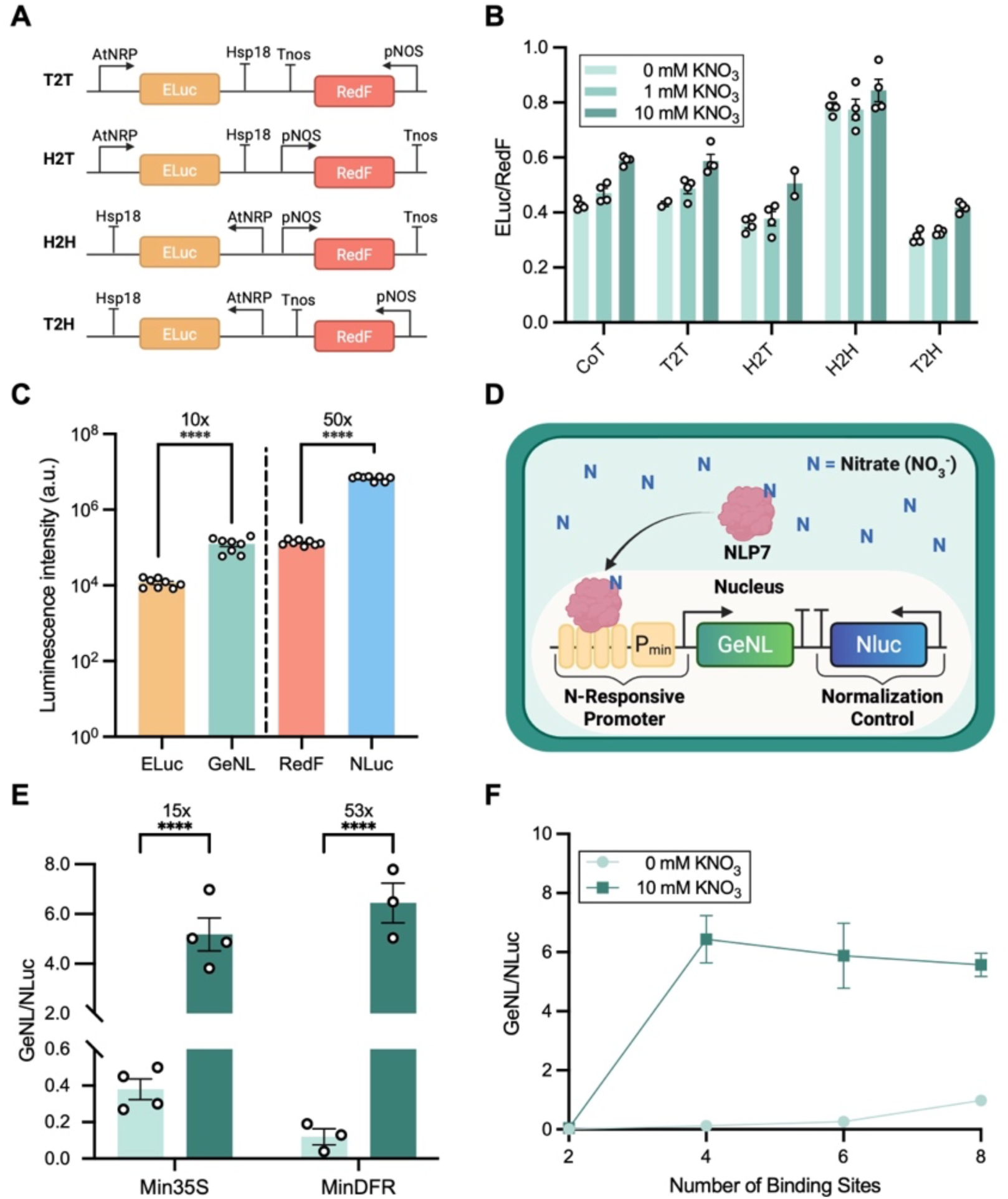
Optimization of the nitrate sensor in *A. thaliana*. **(A)** Schematics of dual luciferase reporter configurations. Head-to-head (H2H), head-to-tail (H2T), tail-to-head (T2H), tail-to-tail (T2T). **(B)** ELuc/RedF signal ratios for different transcriptional unit orientations in *A. thaliana* mesophyll protoplasts under 0, 1, and 10 mM KNO₃. Co-transfection (CoT) of separate ELuc and RedF plasmids was used as a reference condition. Data are shown as mean ± SEM; n = 4 biological replicates with 5 × 10^4^ protoplasts per replicate. **(C)** Luminescence comparison of ELuc/GeNL and RedF/NLuc reporters. Data are shown as mean ± SEM; n = 8 biological replicates with 5 × 10^4^ protoplasts per replicate. Statistical significance was determined using two-tailed unpaired t-tests for each pairwise comparison. **(D)** Schematic of the final NLP7-based nitrate sensor architecture. **(E)** Minimal promoter comparison using min35S and minDFR promoters in intact seedlings. For each promoter, the lighter bars indicate 0 mM nitrate, and the darker bars indicate 10 mM nitrate. Data are shown as mean ± SEM; n = 4 biological replicates with 5 pooled seedlings per replicate. Statistical significance was determined by one-way ANOVA with Tukey’s multiple comparisons test. **(F)** Effect of NLP7 binding site number on sensor output in intact seedlings under 0 and 10 mM nitrate. Data are shown as mean ± SEM; n = 4 biological replicates with 5 pooled seedlings per replicate. ****p < 0.0001.

Although the NRP promoter was suitable for optimizing cassette orientation, it was not used in the final sensor because it responds to multiple nitrogen sources and lacks specificity for nitrate^17^. We therefore developed a new biosensor based on the nitrate-responsive transcription factor NLP7 and its cognate binding sites^25^. The ELuc/RedF reporter pair used for architectural optimization was replaced with a higher-sensitivity GeNL/NLuc luciferase system^26^ to improve signal detection. GeNL generated approximately tenfold higher signal intensity than ELuc, and NLuc produced approximately fiftyfold higher signal intensity than RedF (Fig. 1C). The resulting sensor consisted of tandem NLP7 binding sites positioned upstream of a minimal promoter driving an inducible GeNL reporter, together with a constitutively expressed NLuc normalization cassette arranged in the optimized tail-to-tail configuration (Fig. 1D).

To evaluate sensor performance under more physiologically relevant conditions, we transitioned from protoplast assays to intact *A. thaliana* seedlings grown on MS plates following transient transformation of the sensor circuit^27^. Nitrate responses were quantified as the GeNL/NLuc ratio. We first optimized the minimal promoter downstream of the NLP7 binding sites. Replacing the minimal cauliflower mosaic virus 35S promoter (min35S) with a minimal tomato dihydroflavonol 4 reductase promoter^28^ (minDFR) increased sensor dynamic range from 15-fold to 53-fold between 0 and 10 mM nitrate, reflecting reduced basal activity under nitrate starvation while preserving strong induction (Fig. 1E). The minDFR promoter was therefore used for further optimization.

We next optimized the number of tandem NLP7 binding sites in the synthetic promoter by testing constructs with 2, 4, 6, or 8 binding sites. Sensors containing 2 binding sites did not respond to nitrate, whereas sensors containing 4, 6, or 8 binding sites exhibited 53-fold, 22-fold, and 5-fold induction, respectively, between 0 and 10 mM nitrate (Fig. 1F). Although increasing the number of binding sites enabled nitrate-dependent activation, higher copy numbers also increased basal signal under starvation conditions, resulting in reduced fold change. We therefore selected a final tail-to-tail sensor architecture comprising four tandem NLP7 binding sites upstream of the minDFR promoter and a GeNL/NLuc ratiometric readout.

### Characterization of sensor behavior in plants via transient transformation

Having established the optimized sensor architecture, we characterized its performance in intact *A. thaliana* seedlings following transient transformation^27^ (see Methods). Sensor responsiveness was quantified across nitrate concentrations ranging from 0 to 10 mM, where the sensor exhibited dose-dependent activation that was well described by a saturating Michaelis-Menten relationship (V_max_ = 4.20, K_m_ = 0.57 mM, R^2^ = 0.81; Fig. 2A**; Table S1**). The response was near-linear at low nitrate concentrations between 0 and 1 mM, enabling higher sensitivity under nitrate limitation, and approached saturation above 3 mM nitrate. This profile is consistent with many nitrate-dependent plant transcriptional and physiological responses^29–31^, and indicates that the sensor quantitatively reports nitrate availability across physiologically relevant concentration ranges.

**Figure 2.**
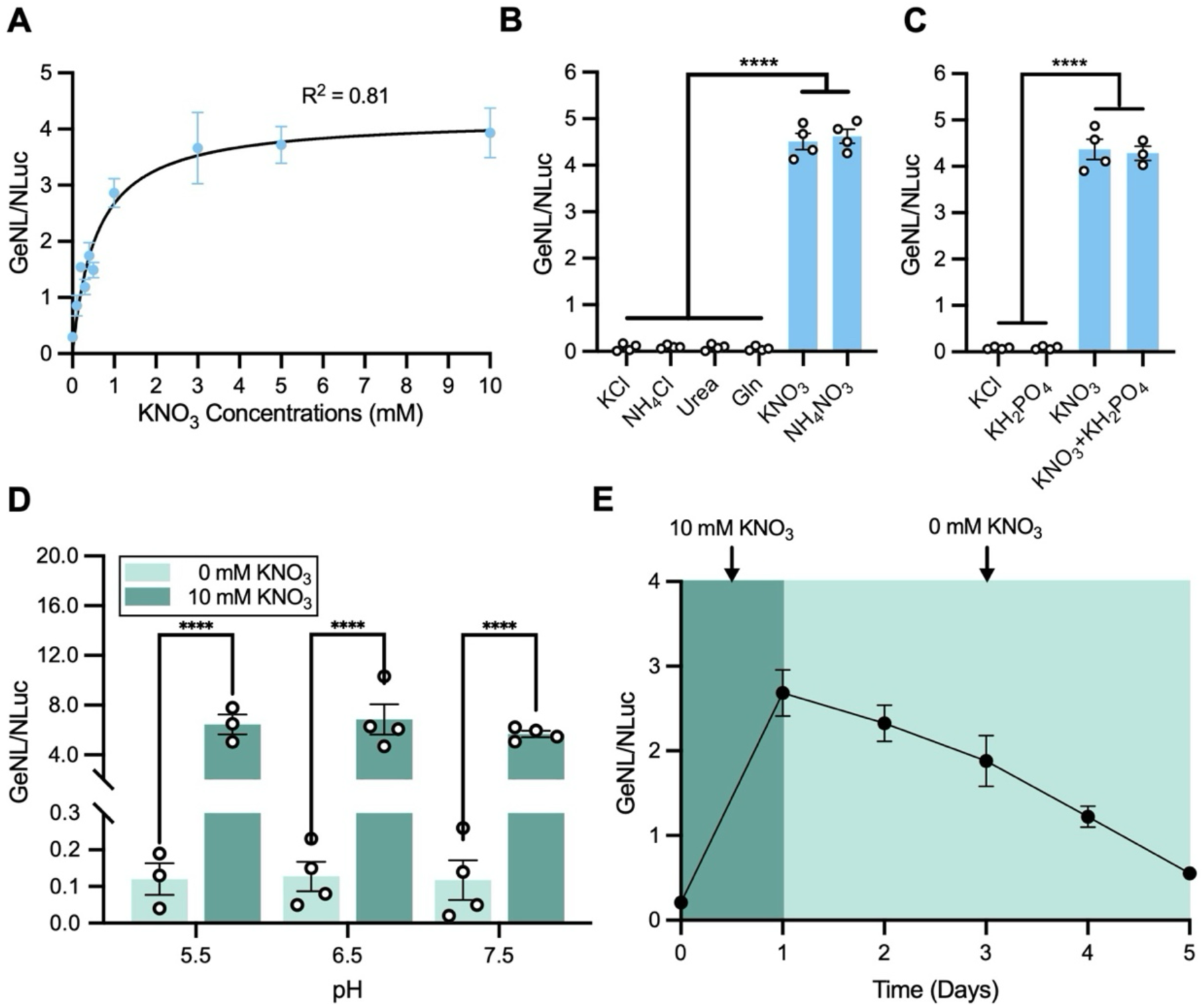
Characterization of sensor behavior via transient expression in Arabidopsis. **(A)** Nitrate dose response curve over 0 to 10 mM nitrate, plotted as GeNL/Nluc and fitted with the Michaelis-Menten equation. **(B)** Nitrogen-source specificity comparing sensor response at 10 mM KCl, NH_4_Cl, urea, glutamine (Gln), KNO_3_, and NH_4_NO_3_. **(C)** Sensor response to phosphate (KH_2_PO_4_) at 1.25 mM, alone or together with 10 mM KNO_3_. **(D)** Sensor output at 0 and 10 mM nitrate conditions at pH 5.5, 6.5, and 7.5. **(E)** Tracking of dynamic sensor response as seedlings were transferred from 10 mM to 0 mM nitrate plates. In all panels, data are shown as mean ± SEM; n = 4 biological replicates with 5 pooled seedlings per replicate. Statistical significance in panels **B** and **C** was assessed using one-way ANOVA with Tukey’s multiple comparisons test, and in panel **D** using Šídák’s multiple comparisons test; ****p < 0.0001.

To determine sensor specificity, we tested alternative nitrogen sources commonly present in soil environments. Potassium nitrate robustly induced the sensor, whereas ammonium, urea, and glutamine did not elicit detectable activation, similar to the potassium chloride negative control (Fig. 2B). Ammonium nitrate activated the sensor to a similar extent as potassium nitrate, indicating that ammonium did not inhibit nitrate-dependent reporter output (Fig. 2B), which was observed in prior nitrate sensors using the NRP promoter^17^. To assess interference from prevalent soil nutrients, we also evaluated the effect of phosphate on sensor activity. Supplementation of high levels of phosphate (1.25 mM) neither triggered sensor activation nor attenuation (Fig. 2C), indicating selectivity for nitrate relative to this major macronutrient.

Sensor robustness was further evaluated across a range of pH values, which often fluctuate under field conditions. From pH 5.5 to 7.5, the sensor maintained strong induction at high nitrate, while preserving low basal output under nitrate-free conditions (Fig. 2D), demonstrating the sensor’s potential to detect nitrate across environmentally relevant pH values. Lastly, to assess whether the sensor responds dynamically and reversibly to changes in nitrate availability, seedlings were first induced with 10 mM nitrate and then transferred to nitrate-free media. Following nitrate removal, sensor output declined toward baseline over 4 days (Fig. 2E), indicating that the sensor reports current nitrate availability rather than retaining a long-term memory of prior exposure.

To evaluate whether this sensing architecture functions beyond Arabidopsis, we tested the sensor in the tomato crop (*Solanum lycopersicum* cv. Micro-Tom). Following transient transformation^27^ (see Methods), tomato seedlings exhibited clear, dose-dependent increases in GeNL/NLuc signal across 0, 1, and 10 mM nitrate treatments **(Supp. Fig. 1)**. These results indicate that the optimized synthetic NLP7-responsive promoter module functions across plant species and suggest that the sensing framework may be transferable to additional crop species.

### Characterization of sensor behavior in *A. thaliana* sentinel plants

To enable long-term monitoring of nitrate dynamics, we generated stable transgenic *A. thaliana* lines expressing the optimized sensor. Three independent homozygous lines were analyzed (**Table S2**). To establish the operational range of the stably integrated sensor, we quantified dose-response relationships for each line between 0 and 10 mM KNO_3_. All three lines displayed dose-dependent activation consistent with saturating Michaelis-Menten kinetics (Fig. 3A**; Table S1)**, with best-fit parameters of V_max_ = 7.27 and K_m_ = 0.12 mM (Line #1), V_max_ = 9.29 and K_m_ = 0.15 mM (Line #2), and V_max_ = 6.49 and K_m_ = 0.08 mM (Line #3). Notably, all stable lines exhibited higher V_max_ values and lower apparent K_m_ values than observed in transient assays, indicating stronger maximal reporter output and increased sensitivity to nitrate following genomic integration. In each case, sensor output was approximately linear at low nitrate concentrations (0-0.5 mM) before approaching saturation at higher concentrations, providing high sensitivity for detecting changes within agronomically relevant low-nitrate ranges.

**Figure 3.**
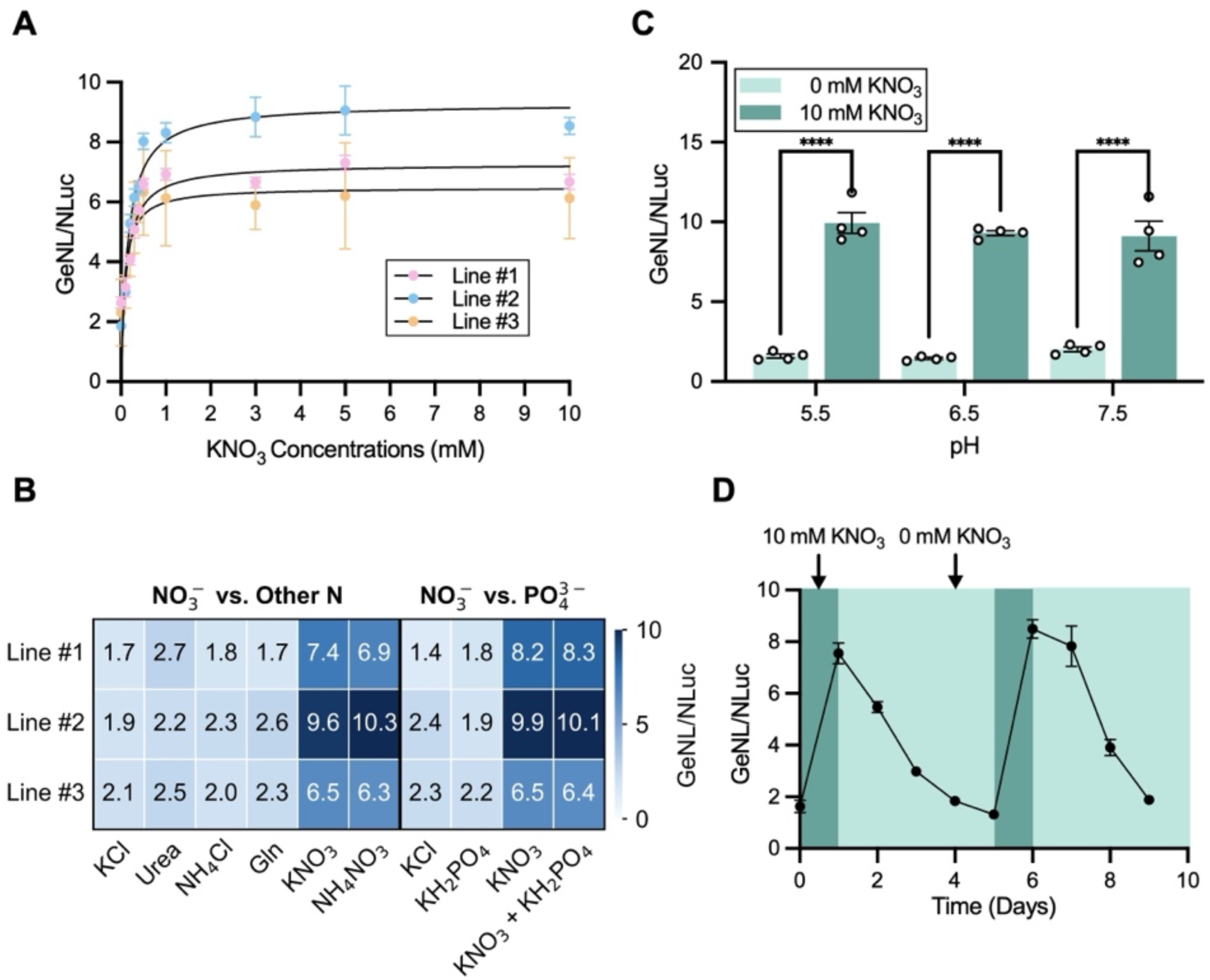
Performance of Arabidopsis sentinel plants as nitrate sensors. **(A)** Nitrate dose response curves of three independent T2 lines (Lines #1 - #3) grown across a KNO₃ gradient (0-10 mM); sensor output fits the Michaelis-Menten kinetics. **(B)** Heatmap showing mean GeNL/NLuc response for each line under varying nitrogen sources and phosphate conditions. **(C)** Sensor output in Line #2 under nitrate-free (0 mM) and nitrate-sufficient (10 mM) conditions at pH 5.5, 6.5, and 7.5. **(D)** Sensor response reversibility in Line #2 during alternating 0 mM and 10 mM KNO₃ treatments; dark green shading indicates 10 mM KNO₃, and light green shading indicates 0 mM KNO₃. In all panels, data are shown as mean ± SEM; n = 4 biological replicates with 5 pooled seedlings per replicate. Statistical significance in panel **C** was assessed using one-way ANOVA with Šídák’s multiple comparisons test; ****p < 0.0001.

We next examined whether genomic integration of the sensor affected its specificity. As in transient assays, robust induction occurred exclusively to nitrate, whereas urea, ammonium, glutamine, potassium chloride, or phosphate did not elicit responses above the baseline levels (Fig. 3B), demonstrating that nitrate selectivity was maintained in the stable sentinel plant lines.

Given its high maximal output, we selected Line #2 for further characterization. Similar to transient assays, sensor performance was maintained across a range of pH values (Fig. 3C). To evaluate response dynamics and reversibility, seedlings from Line #2 were subjected to alternating periods of nitrate supplementation at 10 mM KNO₃ and nitrate withdrawal over a 9-day time-course experiment. Sensor output increased rapidly upon nitrate addition, reached saturation, and then declined toward baseline upon nitrate removal, with this pattern reproducibly observed over two successive cycles (Fig. 3D). These results indicate that the sensor dynamically and reversibly reports changes in nitrate availability rather than exhibiting an irreversible response.

### Sentinel plants report bioavailable nitrate levels in agricultural soils

Defined media offer a controlled way to engineer sensor properties; however, performance in soil is substantially more challenging as real soils comprise heterogeneous physical, chemical, and biological matrices that can interfere with sensor function. We therefore evaluated whether the nitrate-sensing sentinel plants retain quantitative responsiveness in agricultural soils collected from four different California sites with diverse soil properties (Davis, Riverside, Five Points, and Hanford; **Table S3)**. For each soil, nitrate concentrations were quantified using commercial soil analysis performed by Wallace Laboratories (NO_3_^−^ ppm; right y-axis), and sentinel plant output from the same samples was measured as GeNL/NLuc (left y-axis; Fig. 4A). Across field sites, sensor output tracked the analytical nitrate measurements (Spearman ⍴ = 1.0; **Table S4**): Davis soil contained the lowest NO_3_^−^ and produced the weakest sensor response; Riverside and Five Points had intermediate nitrate levels with corresponding sentinel plant signal; and Hanford soil exhibited the highest NO_3_^−^ together with the highest sensor response. Despite the complexities of real agricultural soils, the sensor reliably reported differences in nitrate availability.

**Figure 4.**
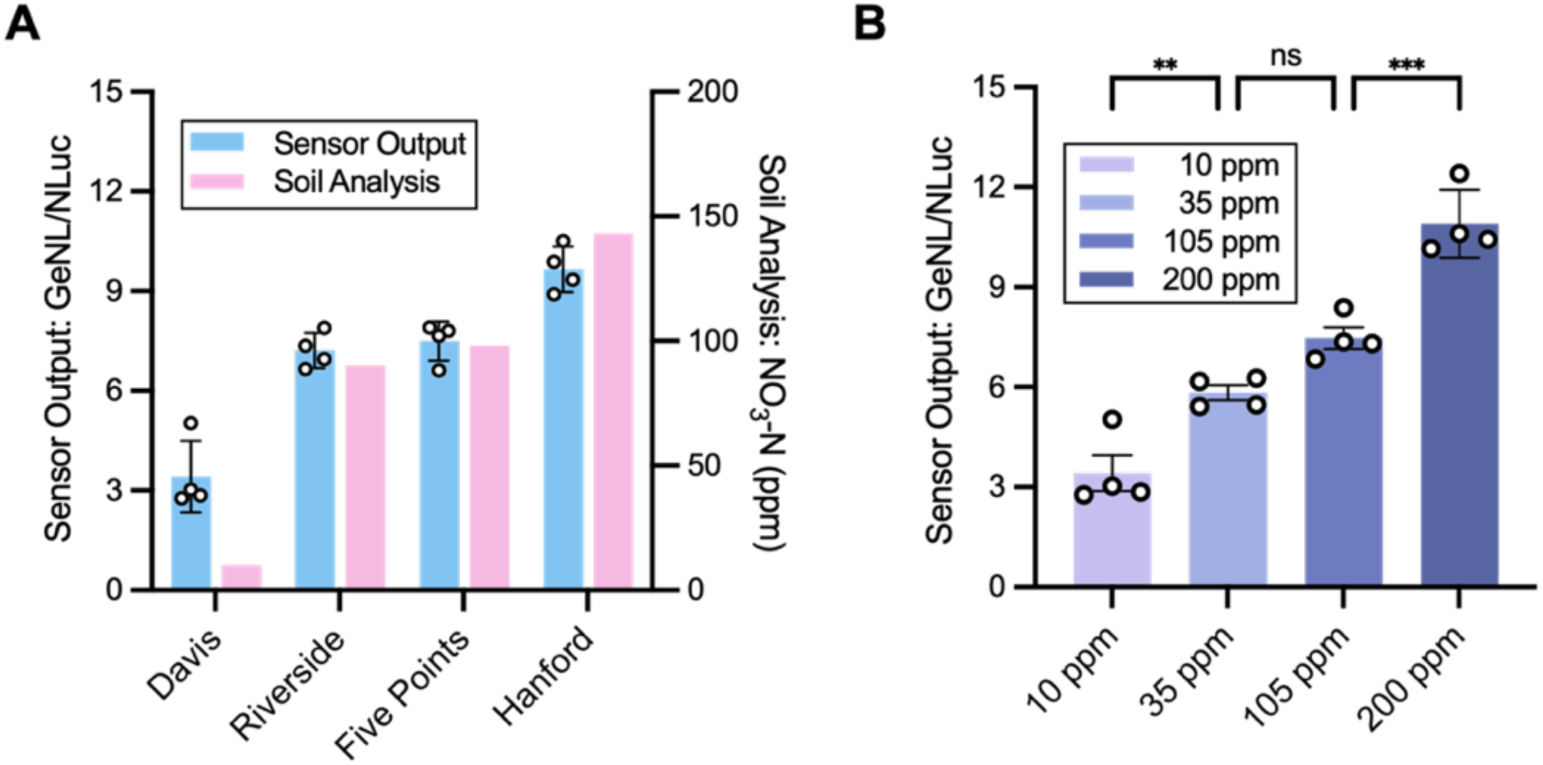
Sentinel plant performance in agricultural soils. **(A)** Sensor output measured in agricultural soils from Davis, Riverside, Five Points, and Hanford compared to quantified nitrate via commercial soil analysis. The left y-axis shows the sentinel plant output as GeNL/NLuc, and the right y-axis shows the ppm level nitrate concentrations measured analytically. **(B)** Sensor response to nitrate supplementation in Davis soil amended with 10, 35, 105, or 200 ppm NO_3_^−^. In both panels, data are mean ± SEM; n = 4 biological replicates with each replicate representing an independent soil sample from the same soil collection batch, in which five seedlings were grown and pooled for luminescence measurements. Statistical analysis was done by one-way ANOVA followed by Tukey’s post hoc test; ns p > 0.05; **p < 0.01; ***p < 0.001.

We next tested whether sentinel plants could resolve incremental nitrate additions within a constant soil background. Using low-nitrate containing Davis soil, we amended the sample with 10, 35, 105, or 200 ppm NO_3_^−^ equivalents and measured the sensor output (Fig. 4B). Sensor output increased in a dose-dependent manner with nitrate supplementation across 10-200 ppm, an agronomically relevant range, indicating that sentinel plants respond quantitatively to increasing nitrate concentrations within agricultural soil matrices.

### Sentinel plants for testing a synthetic nitrogen-fixing consortium in liquid culture

Beyond monitoring nitrate availability in agricultural soils, this genetically encoded sensor can provide a platform for screening synthetic microbial communities for biofertilizer applications. To test whether sentinel plants could monitor microbially mediated nitrogen transformations, we constructed a minimal synthetic microbial consortium designed to convert atmospheric nitrogen into plant-accessible nitrate through sequential nitrogen fixation and nitrification steps. Specifically, we assembled a model synthetic microbiome comprising the diazotroph *Azotobacter vinelandii* (AV), which generates ammonium from atmospheric nitrogen, and the two nitrifying bacteria *Nitrosomonas europaea* and *Nitrobacter winogradskyi*, which convert ammonium into nitrate.

Wild-type *A. vinelandii* (DJ strain) tightly regulates nitrogen fixation and does not typically excrete a significant amount of ammonium into the soil. Therefore, we acquired a mutant *A. vinelandii* strain FM2 (AV FM2) harboring a deletion in the *nifL* regulatory gene, resulting in constitutive nitrogenase activity and ammonium excretion^32^. Consistent with previous reports, when grown in liquid culture, the wild-type strain AV DJ yielded negligible extracellular nitrogen, whereas the mutant AV FM2 strain accumulated approximately 10 and 11 mM ammonium after 4 and 7 days, respectively (Fig. 5A), as quantified using an indophenol colorimetric assay^32^.

**Figure 5.**
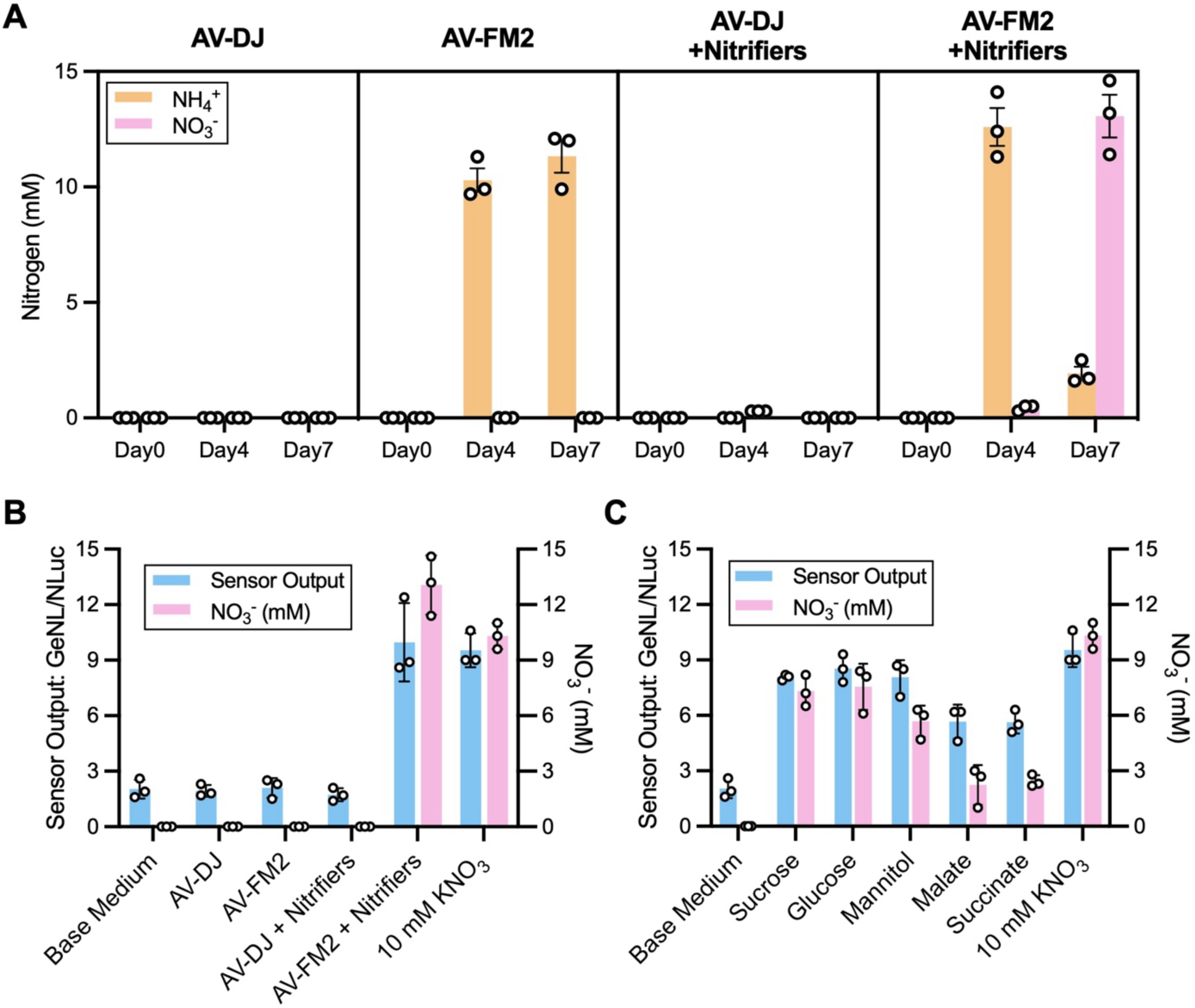
Sentinel plants enable screening of nitrogen-fixing consortia. **(A)** Nitrogen accumulation and transformation in monocultures and co-cultures. Ammonium (NH_4_^+^, yellow bars) and nitrate (NO_3_^-^, pink bars) concentrations were measured over 7 days in wild-type *A. vinelandii* (AV DJ) and the mutant FM2 (AV FM2), with or without the addition of nitrifying bacteria (*N. europaea* and *N. winogradskyi*) at day 4. **(B)** Comparison of sensor output (GeNL/NLuc ratio, blue bars, left y-axis) and chemically measured nitrate concentrations (pink bars, right y-axis) at day 7 for the culture conditions shown in panel A. **(C)** Comparison of sensor output (GeNL/NLuc ratio, blue bars, left y-axis) and chemically measured nitrate concentrations (pink bars, right y-axis) at day 7 for the culture conditions shown in Fig. S2. In all panels, data are mean ± SEM; n = 3 biological replicates, where each replicate represents an independent bacterial culture. For sentinel plant measurements, five seedlings were grown per replicate and pooled for luminescence analysis.

To mimic natural soil nitrification processes, where ammonium is rapidly converted to nitrate, we supplemented the *A. vinelandii* cultures at day 4 with a mixture of nitrifying bacteria (*N. europaea* and *N. winogradskyi*). In cultures containing AV FM2, the accumulated ammonium was rapidly oxidized by the nitrifiers between day 4 and 7, resulting in a final accumulation of ∼13 mM nitrate by day 7, whereas co-cultures containing wild-type AV DJ produced negligible nitrate (Fig. 5A). These ammonium and nitrate levels were quantified using colorimetric assays^32^.

We tested whether sentinel plants could detect these microbially driven nitrogen transformations in a hydroponic setup. Despite high concentrations of ammonium present in the AV FM2 monocultures, the sensor output remained at basal levels, confirming that the system is not activated by ammonium or other microbial metabolites (Fig. 5B). Robust sensor induction was observed only in the AV FM2-nitrifier co-cultures, where the sensor signal matched the 10 mM nitrate positive control, demonstrating high specificity and quantitative response to the final nitrate product even in the presence of other nitrogen species and active microbial populations (Fig. 5B).

Finally, we evaluated whether sentinel plants could be used to screen environmental factors influencing consortium performance, specifically the effect of carbon source. Because nitrogen fixation is energetically demanding, the available carbon source strongly influences nitrogenase activity and ammonium production^33^. We grew the AV FM2-nitrifier co-cultures using five different carbon sources. Cultures grown on sugars or sugar alcohols (sucrose, glucose, and mannitol) accumulated high ammonium concentrations at day 4, which were subsequently converted to 6-8 mM nitrate by day 7. In contrast, cultures grown on organic acids (malate, succinate) supported significantly less nitrogen fixation, resulting in final nitrate concentrations of ∼2 mM, as measured by colorimetric assays **(Supp. Fig. 2)**. When sentinel plants were exposed to these cultures, sensor output quantitatively tracked the differences in nitrate production generated by the various carbon sources, with strong signals in sugar-derived media and lower signals in organic-acid media, consistent with nitrate concentrations measured by chemical assays (Fig. 5C). These results demonstrate that sentinel plants can be used to rapidly screen metabolic outputs of synthetic nitrogen-fixing consortia under varying environmental conditions.

### Testing the synthetic nitrogen-fixing consortium in a model soil via sentinel plants

Ultimately, the practical efficacy of a synthetic nitrogen-fixing consortium depends on its performance in soil rather than liquid culture. To evaluate this, we transitioned to a solid substrate to monitor nitrogen transformations *in situ.* We selected peat moss as a model soil matrix because of its exceptionally low background levels of inorganic nitrogen (**Table S3**), ensuring that detected nitrate would originate from the introduced microbial consortia.

Peat moss pots were inoculated with either engineered AV FM2, wild-type AV DJ, a non-diazotrophic negative-control *Bacillus subtilis*^34^, or a heat-killed biomass negative control with AV FM2 (HK AV FM2). To enable nitrification, *N. europaea* and *N. winogradskyi* were added to all pots on day 7, and nitrogen dynamics were monitored over a 13-day period. The negative control conditions (heat-killed AV FM2 and *B. subtilis*) showed negligible nitrogen accumulation throughout the experiment, as measured by both colorimetric assays and sentinel plant outputs **(Supp.** Fig. 3; Fig. 6A, B). By day 7, prior to the addition of nitrifiers, both *Azotobacter* strains had enriched peat moss with ammonium, with the AV FM2 producing significantly more ammonium (∼220 ppm) than the wild-type AV DJ (∼170 ppm) **(Supp. Fig. 3).**

**Figure 6.**
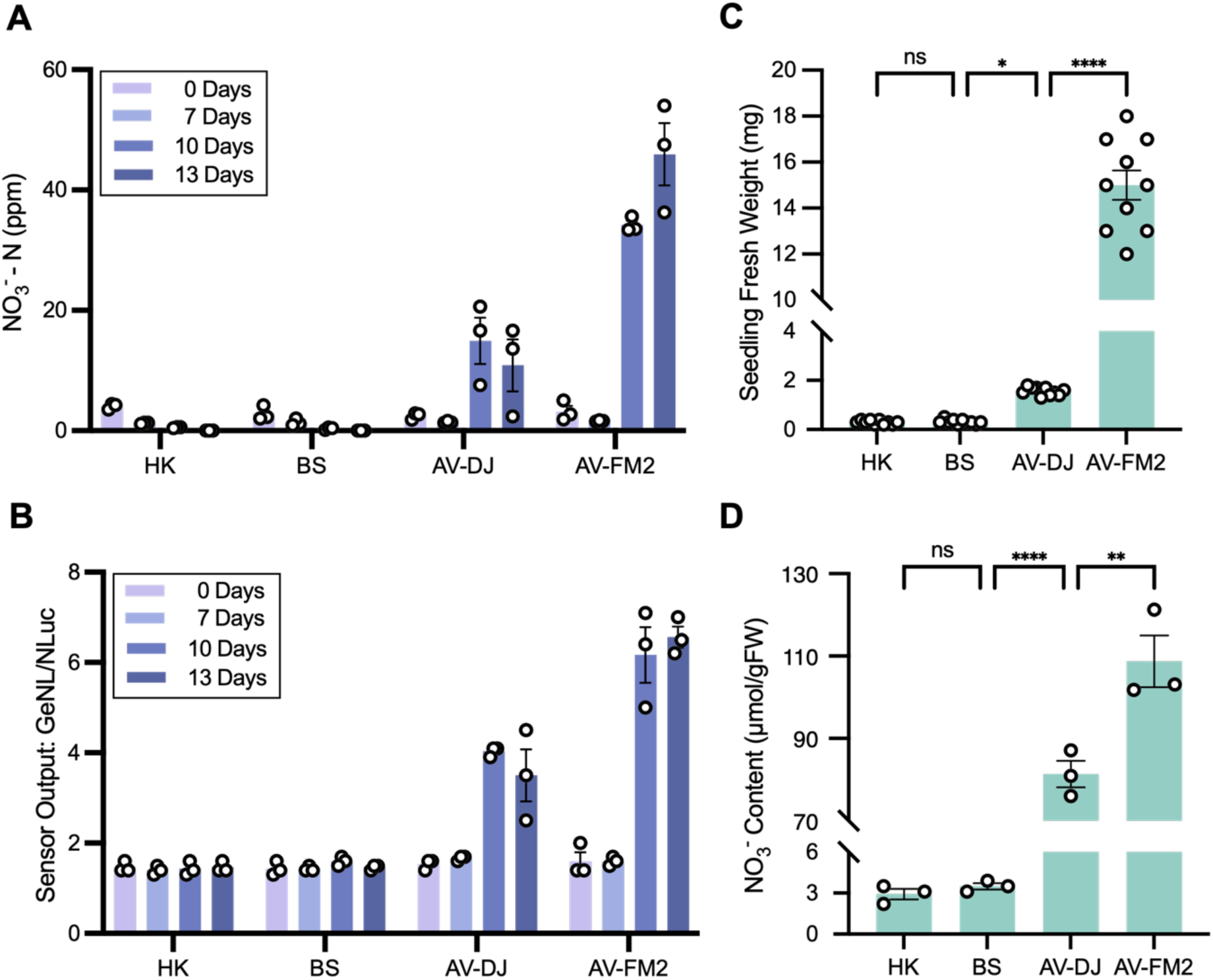
Monitoring nitrogen transformations via sentinel plants and plant growth promotion. **(A)** Nitrate accumulation in peat moss inoculated with bacterial treatments over a 13-day period. Nitrifiers were introduced to all pots at day 7. HK: heat-killed *AV* FM2 biomass control; BS: *B. subtilis* non-diazotrophic control; AV DJ: wild-type *A. vinelandii*; AV FM2: engineered mutant. **(B)** *In situ* sensor performance within the peat moss matrix treated with various bacterial treatments over a 13-day period. Nitrifiers were introduced to all pots at day 7. In panels **A** and **B**, n = 3 biological replicates, each representing an independent culture. For sentinel measurements in panel **B**, five seedlings were grown per replicate and pooled for luminescence analysis. **(C)** Fresh weight of *A. thaliana* seedlings was assessed 14 days post-germination. n = 10 seedlings, with each seedling representing one replicate. **(D)** Internal tissue nitrate content (μmol/g FW) of *A. thaliana* seedlings was assessed 14 days post-germination. Data represent mean ± SEM; n = 3 biological replicates, each consisting of 10 pooled seedlings. Statistical significance was determined by one-way ANOVA with Tukey’s post-hoc test; ns p > 0.05; *p < 0.05; **p < 0.01; ****p < 0.0001.

Following nitrifier addition, this ammonium pool was progressively converted to nitrate. The higher ammonium production by AV FM2 resulted in significantly greater soil nitrate accumulation, reaching approximately 35 ppm by day 10 and 45 ppm by day 13 (Fig. 6A). In contrast, pots inoculated with AV DJ accumulated substantially less nitrate, plateauing at approximately 10-14 ppm during the same period (Fig. 6A). To monitor these nitrogen transformations *in situ*, sentinel plants were introduced into the inoculated peat moss soil. Sensor output closely tracked nitrate formation, producing strong signals in AV FM2 pots at days 10 and 13 that corresponded to the elevated soil nitrate concentrations (Fig. 6B). Sensor responses in AV DJ pots were substantially lower, reflecting the reduced nitrate accumulation, whereas HK AV FM2 and *B. subtilis* control pots remained at baseline sensor output levels.

To demonstrate the establishment and persistence of the microbial consortium, we performed 16S rRNA gene amplicon sequencing on peat samples collected at 0, 7, 10, and 13 days from both the AV DJ and AV FM2 treatments (**Supp.** Fig. 4). Community profiling confirmed the establishment of *Azotobacter* in all inoculated pots. Following nitrifier addition, reads corresponding to *Nitrosomonas* and *Nitrobacter* increased in abundance, coinciding with nitrate accumulation. Together, these data indicate that the observed ammonium and nitrate pools arise from the activity of the introduced diazotroph-nitrifier consortium and that sentinel plants accurately report these transformations *in situ* in soil.

To evaluate the biofertilizer potential of this three-member synthetic consortium, wild-type *Arabidopsis thaliana* plants were grown in treated peat moss pots, and plant growth was assessed after 14 days. Nitrate produced by the AV FM2-nitrifier consortium supported robust plant growth **(Supp.** Fig. 5), yielding seedlings with a fresh weight of 15 mg each, compared with 1.5 mg each in the AV DJ-nitrifier treatment and substantially lower biomass in the heat-killed AV FM2 and *B. subtilis* control (Fig. 6C).

Analysis of these plant tissues confirmed that enhanced growth was associated with increased nitrogen uptake. AV FM2-nitrifier-treated seedlings accumulated significantly higher internal nitrate levels (110 μmol g^-1^ fresh weight) than AV DJ-nitrifier-treated seedlings (80 μmol g^-1^ fresh weight), as measured by a colorimetric assay (Fig. 6D). Relative to the heat-killed AV FM2 and *B. subtilis* control, these values correspond to approximately 37-fold and 28-fold increases in internal nitrate content for AV FM2 and AV DJ treatments, respectively. Together, these results demonstrate that the synthetic consortium promotes plant growth in a soil-like environment through a controlled nitrogen-fixation and nitrification pathway that can be accurately and quantitatively monitored by sentinel plants.

## Discussion

This study establishes that plants can function as integrated environmental nitrate sensors that directly report plant-accessible nutrient availability. We developed a genetically encoded plant biosensing system that enables quantitative monitoring of plant-accessible nitrate across controlled media, agricultural soils, and microbial ecosystems. Because the reporter operates downstream of endogenous nitrate-sensing pathways, the resulting signal reflects nitrate availability experienced by the plant rather than concentrations obtained from soil extraction assays. This distinction is important because plant nutrient status is shaped not only by bulk soil chemistry but also by root uptake, transport, and local microenvironmental conditions. Sentinel plants, therefore, provide a plant-integrated measurement of nitrate availability that complements analytical measurements of soil chemistry and existing biosensors.

The sensing architecture combines a synthetic NLP7-responsive promoter with a ratiometric luciferase reporter system optimized for whole-plant measurements. Systematic optimization of promoter configuration, binding-site number, and reporter architecture revealed design principles for quantitative plant biosensing. Transcriptional orientation of reporter cassettes strongly affected signal fidelity, with the tail-to-tail configuration minimizing transcriptional interference while preserving inducible behavior. This result is consistent with prior studies of compositional context in synthetic gene circuits^23,35^ and highlights circuit architecture as an important design variable in plant synthetic biology. Tuning the number of NLP7 binding sites further demonstrated that promoter architecture governs dynamic range and basal activity, with four binding sites providing the optimal balance between strong activation and low background.

Prior plant nitrate sensors primarily report intracellular nitrate dynamics rather than environmental nutrient availability. In contrast, the sentinel plant system described here reports nitrate availability integrated through plant uptake. Functional characterization showed that the sensor exhibits a dose-dependent response approximating Michaelis-Menten kinetics, consistent with prior findings of nitrate-responsive transcriptional programs in plants^29^. The sensor exhibits strong specificity for nitrate over ammonium, urea, glutamine, and phosphate, thereby avoiding the issues observed in some previous nitrate-responsive promoters, in which ammonium suppressed the response^17^. Dynamic withdrawal experiments further showed that the reporter reversibly tracks changes in nitrate availability rather than maintaining an extended transcriptional memory of past exposure. These properties are important for environmental monitoring, where nutrient levels fluctuate over time and selectivity and reversibility are required for meaningful interpretation.

Stable genomic integration preserved the performance across Arabidopsis lines, demonstrating that the sensing architecture functions reliably beyond transient assays. Although absolute signal amplitudes varied among lines, likely due to positional effects associated with transgene insertion, response kinetics and nitrate specificity were consistent. These results indicate that the sensing architecture is robust to genomic context, enabling long-term monitoring of nitrate dynamics in stable plant material. Importantly, transient experiments in tomato demonstrated that the sensing architecture also functions in a crop species, indicating that the NLP7-based promoter module and ratiometric reporter design are transferable across species. This cross-species functionality suggests that similar sensing systems could be implemented in agriculturally relevant crops.

A key step toward practical relevance is demonstrating that plant biosensors remain quantitative in complex soil matrices. Across agricultural soils collected from multiple California field sites, sentinel plant output closely tracked nitrate levels measured using standard analytical methods. The sensor also resolved incremental nitrate additions within a constant soil background across agronomically relevant concentration ranges. These results indicate that plant-based biosensors can operate reliably despite the physical, chemical, and biological heterogeneity inherent to soil environments, demonstrating that engineered plants can potentially report nutrient variation under realistic environmental conditions.

An important application of this sensing platform is the ability to monitor microbial nitrogen transformations that generate plant-accessible nitrate. Using sentinel plants as readouts, we identified a minimal three-member microbial consortium capable of converting atmospheric nitrogen into plant-available nitrate through sequential nitrogen fixation and nitrification. Sentinel plant output quantitatively tracked nitrate generated by this microbial community in both liquid culture and soil-like matrices, demonstrating that plant-based biosensors can report microbial metabolic activity relevant to plant nutrition. The current setup used a low-background peat substrate to isolate consortium-driven nitrogen transformations under controlled conditions, and future studies should evaluate sensor performance and consortium function in agricultural soils to assess robustness under realistic environmental complexity.

This consortium represents a potential nitrogen biofertilizer strategy, as its activity increased nitrate availability, tissue nitrate accumulation, and plant biomass. Nitrate concentrations reached 45 ppm by day 13, which falls within agronomically relevant nitrate levels^36–38^. At the same time, nitrate is highly mobile in soil, susceptible to leaching, and nitrification processes can contribute to nitrous oxide emissions. Future implementation can incorporate strategies that synchronize nitrate generation with plant demand while minimizing environmental nitrogen loss. This platform also enabled rapid evaluation of environmental factors that influence nitrogen fixation efficiency. Variation in carbon source availability substantially altered ammonium production by *Azotobacter vinelandii*, which in turn influenced downstream nitrate accumulation following nitrification. Carbon sources such as sucrose, glucose, and mannitol supported higher nitrate production than organic acids such as malate and succinate, consistent with previous reports that carbon metabolism strongly regulates diazotrophic nitrogen fixation^33^. Sentinel plant output accurately reflected these shifts in nitrate production, illustrating how plant-integrated sensing platforms can accelerate design-build-test cycles for optimizing microbial communities.

Extending these capabilities toward agricultural deployment will require additional engineering of the platform. Currently, the sensor relies on a luciferase-based readout that requires exogenous substrate addition, which could be sprayed for *in situ* detection. However, future iterations should incorporate autonomous bioluminescence pathways^39,40^ or replace luminescence with visible reporters, such as pigment-based reporters (e.g., RUBY^41^) or UV-excitable fluorescent proteins^42^, enabling direct visualization and compatibility with camera- or drone-based imaging for *in situ* large-scale monitoring. Future work should also evaluate how diurnal transcriptional dynamics^43^ and nitrate interactions with other nutrients^44^ influence sensor performance under field conditions and account for these effects.

More broadly, the sensing architecture developed here establishes a framework for engineering plants as environmental biosensors that can monitor diverse soil and rhizosphere processes in agricultural ecosystems. Because the platform relies on modular promoter design and ratiometric normalization, similar approaches could be adapted to detect additional nutrients, metabolites, or environmental cues relevant to plant growth. These sensing modules could also be coupled to regulatory circuits that alter plant behavior in response to nutrient status, extending sentinel plants from passive reporters toward responsive plants with programmable environmental adaptation.

## Methods

### DNA Constructs

Constructs were designed using SnapGene software (SnapGene, GSL Biotech). All plasmids were assembled through BsaI/SapI-mediated Golden Gate loop assembly. A comprehensive list of standard parts and assembled expression constructs is provided in **Supplementary Table 5**. Plasmids and sequence information are available through Addgene (**Table S5** for Addgene IDs).

### Bacterial Strains and Growth Conditions

*Agrobacterium tumefaciens* strain GV3101 (1282PS-12, Intact Genomics) was obtained from Intact Genomics and routinely cultured in LB medium. *Azotobacter vinelandii* strains DJ and FM2 were obtained as gifts from the Ané Laboratory (University of Wisconsin-Madison) and routinely cultured in Burk’s medium^45^. *Nitrosomonas europaea* (19718, ATCC) was obtained from the American Type Culture Collection (ATCC) and routinely cultured in ATCC Medium 2265. *Nitrobacter winogradskyi* (25391, ATCC) was obtained from ATCC and routinely cultured in DSMZ Medium 756a. Unless otherwise stated, all strains were incubated at 30°C with shaking at 250 rpm.

### Seed Sterilization and Germination

*A. thaliana* (Col-0) seeds were surface-sterilized by exposure to chlorine gas. Chlorine gas was generated by combining 100 mL of bleach with 3 mL of 6 N hydrochloric acid. The mixture was placed in a desiccator inside a chemical fume hood, and the seeds were exposed to the resulting gas for at least 3 hours. After sterilization, seeds were either stored or used immediately, with sterile conditions maintained throughout. *Solanum lycopersicum* (tomato, cultivar Micro-Tom) seeds were surface-sterilized in 33% (v/v) bleach containing 0.1% Tween-20 for 6 minutes. Following sterilization, seeds were rinsed four to five times with sterile water. All seeds were germinated on ½-strength MS plates containing 0.5× Murashige and Skoog Basal Salt Mixture (M524, PhytoTech Labs), 0.05% (w/v) MES, and 0.7% Phytagel (P8169, Millipore Sigma), adjusted to pH 5.7. Plates were incubated in a growth chamber at 22°C under a 16/8 h light/dark cycle.

### Isolation and Transformation of *A. thaliana* Protoplasts

Mesophyll protoplasts were isolated and transformed from fully expanded leaves of 4-week-old *A. thaliana* (Col-0) following the protocol of Muchenje et al^46^. After transformation with the nitrate-responsive constructs, protoplasts were incubated for 24 h in WI solution supplemented with the specified NO_3_^-^ concentrations. Afterward, luminescence was measured using a TECAN plate reader to quantify sensor output, as described in the *Quantification of Sensor Output* section.

### Transient Transformation of *A. thaliana* Seedlings

Nitrate-responsive constructs were transiently transformed into 7-day-old *A. thaliana* seedlings using the VAST (Vacuum- and Sonication-Assisted Transient Expression) protocol^27^. After 2 days of co-cultivation with *A. tumefaciens*, seedlings were thoroughly rinsed with sterile water and transferred to nitrogen-free ½ MS plates (M531 lacking nitrogen, PhytoTech Labs) supplemented with 10 mM KCl. Seedlings were maintained under nitrogen-depleted conditions for 3 days to re-establish basal sensor output.

Following nitrogen starvation, seedlings were transferred to nitrogen-free ½ MS plates containing 100 µM timentin and the indicated treatments: defined concentrations of KNO₃ or 10 mM of alternative nitrogen sources, with supplemental KCl added as required to ensure equivalent potassium levels across treatments. Additional assays were performed on plates buffered to specified pH values. Treatments were applied for 24 hours, after which luminescence was quantified to assess sensor output following the protocol described below.

### Transient Transformation of *Solanum lycopersicum* (cv. Micro-Tom) Seedlings

Transient transformation of Micro-Tom was carried out using the VAST protocol^27^. Seeds of *S. lycopersicum* (cv. Micro-Tom) were germinated on nitrogen-free ½ MS plates (M531 lacking nitrogen, PhytoTech Labs) until cotyledons were fully expanded (approximately 9 days). Seedlings were then co-cultivated with *A. tumefaciens* carrying the sensor plasmid for 2 days. Following co-cultivation, seedlings were transferred to nitrogen-free ½ MS plates supplemented with 100 µM timentin and defined concentrations of KNO_3_. Treatments were applied for 24 hours, after which luminescence was quantified to assess sensor output following the protocol described below.

### Preparation of agricultural soil

Agricultural soils collected from California field sites were passed through a 2-mm sieve to remove rocks and large organic debris. To improve root-zone aeration and maintain a consistent physical structure after debris removal, the sieved soil was mixed with an equal mass of peat moss. For experiments, soil moisture was maintained at 70% field capacity, defined as 70% of the water content retained by saturated soil after gravitational drainage. Field capacity was determined gravimetrically by measuring the difference between the weight of saturated soil after drainage and the dry soil weight, and experimental soils were adjusted to 70% of this value prior to inoculation or planting.

### Preparation and Application of Sentinel Plants

Transformation of *A. thaliana* was performed by spraying liquid suspension of *A. tumefaciens* GV3101 carrying the sensor constructs onto plant inflorescences and selection of transgenic T1 and T2 seeds using a fluorescence microscope to visualize the seed-specific FAST-red marker. Transgene presence in homozygous lines was confirmed with conventional PCR and subsequent electrophoresis. Homozygous *A. thaliana* lines carrying the sensor constructs were germinated on standard ½ MS medium (M524, PhytoTech Labs) for 9 days. Seedlings were then transferred to nitrogen-free ½ MS plates (M531 lacking nitrogen, PhytoTech Labs) supplemented with 10 mM KCl and maintained under nitrogen-depleted conditions for 3 days to re-establish basal sensor output.

Following nitrogen starvation, seedlings were transferred to nitrogen-free ½ MS plates containing the indicated treatments: defined concentrations of KNO₃ or 10 mM of alternative nitrogen sources, with supplemental KCl added as required to ensure equivalent potassium levels across treatments. Additional assays were performed on plates buffered to specified pH values. Treatments were applied for 24 hours, after which luminescence was quantified to assess sensor output following the protocol described below.

In soil-based assays, sensor performance was evaluated using a soil-peat moss mixture prepared by combining sieved California agricultural soil and peat moss in a 1:1 (w/w) ratio. Seedlings were incubated in the indicated treatments for 24 hours, after which luminescence was quantified using the same protocol.

### Quantification of Sensor Output

Luminescence signals were quantified by collecting leaves from five seedlings or 5 × 10^5^ protoplasts per sample into 2-mL tubes (15-340-162, Fisher Scientific) preloaded with 2.8-mm ceramic beads (15-340-160, Fisher Scientific). In all experiments, samples were collected in the afternoon to reduce the impact of diurnal effects, immediately flash-frozen in liquid nitrogen, and homogenized using a minibead-beater tissue disruptor (15-340-164, Fisher Scientific) for 40 seconds at maximum speed. Subsequently, 100 µL of 1× Cell Lysis Buffer (E1531, Promega) was added to each tube, and samples were vortexed vigorously until no visible particulates remained. Lysates were placed on ice for 10 minutes and centrifuged at 5,000g for 5 minutes to pellet debris.

Luminescence was measured using the Nano-Glo Luciferase Assay System (N1110, Promega) according to the manufacturer’s instructions with minor modifications. For each reaction, 15 µL of clarified protein extract was mixed with 30 µL of Nano-Glo reagent in a 96-well plate (07-200-627, Fisher Scientific). Luminescence was recorded on a Tecan Spark plate reader at 25°C using GeNL (520-590 nm) and NLuc (430-485 nm) filters with an integration time of 100 ms per well. Total, GeNL, and NLuc signals were acquired, and spectral deconvolution was performed using the Promega ChromaLuc Technology Calculator (Technical Manual TM062).

### Nitrate-Forming Medium and Liquid Co-culture Conditions

The nitrate-forming medium (NFM) used in this study consisted of a base medium supplemented with a sterile 100× phosphate buffer. The base medium was prepared by dissolving 153 mg of M407 (M407, PhytoTech Labs) and 10 g of sucrose in ultrapure water and adjusting the volume to 500 mL. The 100× phosphate buffer was prepared by dissolving 1.10 g of KH₂PO₄, 5.77 g of K₂HPO₄·3H₂O, and 7.5 g of KHCO₃ in ultrapure water and adjusting the final volume to 50 mL. The base medium was sterilized by autoclaving, and the phosphate buffer was sterilized by filtration and added to the base medium at a 1:99 ratio (e.g., 100 μL per 9.9 mL of medium).

Co-cultures of *A. vinelandii* (strains DJ or FM2), *N. europaea* (ATCC 19718), and *N. winogradskyi* (ATCC 25391) were established by first cultivating each organism independently under its routine culture conditions. *A. vinelandii* was inoculated into freshly prepared NFM at an initial optical density of OD₆₀₀ = 0.01 and cultured for 4 days. Subsequently, *N. europaea* and *N. winogradskyi* were introduced by harvesting volumes of each culture equal to the volume of the *A. vinelandii* culture (e.g., 10 mL NE and 10 mL NW for a 10 mL AV culture), centrifuging to collect the cell pellets, and resuspending the pellets directly into the *A. vinelandii* culture. No additional medium was added during this step. All co-cultures were incubated at 30°C with shaking at 250 rpm unless otherwise specified.

### Co-cultivation of Bacteria in Peat Moss

Co-cultures of *Azotobacter vinelandii* (strains DJ or FM2), *Nitrosomonas europaea* (ATCC 19718), and *Nitrobacter winogradskyi* (ATCC 25391) were established by first culturing each organism independently under its respective standard growth conditions. 25g of peat moss was weighed and placed into a Magenta box (30930007, Magenta), and the pH was adjusted to approximately 7.5 using KOH.

Overnight cultures of *A. vinelandii* were harvested by centrifugation and inoculated into freshly prepared NFM at an OD_600_ calculated to achieve a final concentration of approximately 1 × 10⁸ A*. vinelandii* cells per gram of peat moss when applied at 70% field capacity. The peat moss and bacteria were thoroughly mixed by shaking and stirring to ensure uniform distribution. The container was sealed with Micropore tape (19571, Qiagen) to minimize water loss while allowing gas exchange and incubated in a plant growth chamber at 22 °C under a 16/8-h light/dark cycle.

Moisture levels were monitored weekly by measuring water loss, and NFM was added as needed to maintain 70% field capacity. On day 7, *N. europaea* and *N. winogradskyi* were introduced by harvesting 55 mL of each culture, centrifuging to collect cell pellets, and resuspending the pellets directly into the NFM used for moisture maintenance. The peat moss was then thoroughly mixed to distribute the nitrifying bacteria and resealed with Micropore tape. Ammonium and nitrate concentrations in the peat moss system were quantified by colorimetric assays on days 7, 10, and 13 to monitor nitrogen transformation dynamics. In parallel, peat moss samples were collected at the same time points for 16S rRNA gene sequencing to assess microbial community composition.

### Ammonium Quantification

Ammonium concentrations were quantified using a colorimetric assay based on the Berthelot reaction, adapted from Mus et al^32^. Samples and NH₄Cl standards (0-5 mM) were prepared, and 50 µL aliquots were reacted with 100 µL of phenol-nitroprusside reagent (composition: 1% [w/v] phenol and 0.005% [w/v] sodium nitroprusside), followed by the addition of 100 µL of alkaline hypochlorite solution (composition: 1% [v/v] sodium hypochlorite and 0.5% [w/v] NaOH). Reactions were incubated in 1.5 mL Eppendorf tubes at 37 °C for 30 min, after which absorbance was measured at 620 nm. Ammonium concentrations were determined by interpolation from a standard curve. Measurements were performed with at least two technical replicates and three biological replicates, unless otherwise noted.

### Nitrate and Nitrite Quantification

Nitrate and nitrite concentrations were quantified using the Griess assay adapted from Hachiya et al.^47^ Standards (0-320 µM) were prepared using either nitrate or nitrite, as appropriate. For nitrate quantification, 50 µL of sample or standard was mixed with 50 µL of sulfanilamide–VCl₃ reagent (0.25% [w/v] VCl₃ and 1% [w/v] sulfanilamide in 1 M HCl), followed by the addition of 50 µL of N-(1-naphthyl) ethylenediamine solution (0.02% [w/v] in water). Reactions were incubated at room temperature for 6–8 h prior to measurement. For nitrite quantification, VCl₃ was omitted from the sulfanilamide reagent, nitrite standards were used, and reactions were incubated at room temperature for 10 min. Absorbance was measured at 540 nm, and nitrate or nitrite concentrations were determined by interpolation from the corresponding standard curve. All measurements were performed with at least two technical replicates and a minimum of three biological replicates, unless otherwise indicated.

### 16S rRNA Gene Amplicon Sequencing and Microbial Community Analysis

Microbial community composition was assessed using full-length 16S rRNA gene amplicon sequencing. DNA was extracted from peat moss samples using the DNeasy PowerSoil Pro Kit (47014, Qiagen) according to the manufacturer’s instructions. Extracted DNA was used as a template for amplification of the full-length 16S rRNA gene (∼1.5 kb) using the Oxford Nanopore Microbial Amplicon Barcoding Kit 24 (SQK-MAB114.24) following the manufacturer’s protocol (Oxford Nanopore Technologies). Briefly, approximately 10 ng of genomic DNA per sample was used as input for PCR amplification with kit-provided primers targeting the bacterial 16S rRNA gene. Amplicons were barcoded during PCR to enable multiplexing of samples.

Barcoded PCR products were pooled and purified using magnetic bead cleanup prior to adapter attachment. Libraries were prepared according to the Oxford Nanopore microbial amplicon sequencing workflow and loaded onto R10.4.1 flow cells (FLO-MIN114) for sequencing on a Nanopore platform. The analysis pipeline was adapted from Aja-Macaya et al.^48^ In summary, reads were first base-called using the Dorado basecaller super accurate (SUP) model, then demultiplexed with Dorado. The resulting reads were quality-controlled using chopper, filtered for a minimum quality score of 20, a minimum length of 1200, and a maximum length of 1900, and trimmed by 20 nt at the front and back of the sequence. Reads were then aligned and identified with Emu using the default database^48^.

### Nitrogen Concentration Calculations

Nitrogen concentrations were reported using different units depending on the experimental matrix. In liquid culture, nitrogen concentrations were expressed in millimolar (mM), representing millimoles of nitrogen per liter of solution. In soil samples, nitrogen concentrations were expressed in parts per million (ppm), equivalent to mg N kg^-1^ dry soil. Ammonium and nitrate concentrations are reported as nitrogen equivalents, specifically NH_4_^+^–N and NO_3_^-^–N. Thus, reported values represent the nitrogen derived from ammonium or nitrate rather than the total mass of the ionic species.

## Data, Materials, and Software Availability

All data supporting the findings of this study are included in the paper and its Supplementary Information files or are available from the corresponding author upon request. Plasmids generated in this study have been deposited in Addgene **(Table S5)**. Stable transgenic sensor plant lines described in this study have been deposited at the Arabidopsis Biological Resource Center (ABRC) and are available under stock numbers [will be added once paper is accepted]. The 16S rRNA gene sequencing data have been deposited in NCBI under BioProject accession number PRJNA1438707. Data analysis was performed using the code in Aja-Macaya et al^48^.

## Funding

This research was supported by grants from the USDA NIFA Agriculture and Food Research Initiative (Award Number: 2024-67019-42486), Henry Luce Foundation, Shurl Kay Curci Foundation, NASA Translational Research Institute for Space Health (TRISH) through Caltech Space-Health Innovation Fund, and Caltech internal seed and startup funds. EL is supported by the National Science Foundation (NSF) Graduate Research Fellowships Program (GRFP). CB is supported through the NSF GRFP and Caltech’s Biotechnology Leadership Pre-Doctoral Training Program. TMO is partially supported by the U.S. NSF under Cooperative Agreement No. 2330145. Any opinions, findings, and conclusions or recommendations expressed in this material are those of the author and do not necessarily reflect the views of the U.S. NSF.

## Author contributions

E.L. and G.S.D. conceived the project, designed the study, and wrote the manuscript. E.L. performed most experiments and conducted all data analysis. C.B. performed 16S sequencing and optimized growth conditions for ammonia-fixing and nitrifying bacteria in peat moss. T.O. generated stable sentinel plant lines. E.G., C.G., Y.W., and J.J. performed colorimetric assays for nitrogen compounds and optimized growth conditions for ammonia-fixing and nitrifying bacteria in liquid culture.

## Acknowledgements

We would like to thank Nicola Patron’s lab for providing the NLP7 binding site sequences. We thank Julie Marie Pelletier for coordinating and shipping soil samples from Davis, Steven Ries and Peggy A. Mauk for assisting us in collecting soil samples from Riverside and introducing the characteristics of the soil, and Lauren Hale for providing and shipping soil samples from Five Points and Hanford. We also thank Jean-Michel Ané for providing the *A. vinelandii* FM2 strain. We would also like to thank Prof. Hikmet Geckil for his assistance in editing the manuscript.

## Competing interests

The authors declare that they have no known competing financial interests or personal relationships that could have appeared to influence the work reported in this paper.

## Notes

### Competing Interest Statement

The authors have declared no competing interest.

## References

1. Robertson, G. P. & Vitousek, P. M. Nitrogen in agriculture: balancing the cost of an essential resource. Annu. Rev. Environ. Resour. 34, 97–125 (2009).

2. Jones, M. N. Nitrate reduction by shaking with cadmium: alternative to cadmium columns. Water Res. 18, 643–646 (1984).

3. Dick, W. A. & Tabatabai, M. A. Ion chromatographic determination of sulfate and nitrate in soils. Soil Sci. Soc. Am. J. 43, 899–904 (1979).

4. Maggini, R., Carmassi, G., Incrocci, L. & Pardossi, A. Evaluation of quick test kits for the determination of nitrate, ammonium and phosphate in soil and in hydroponic nutrient solutions. Agrochimica 54, 331–341 (2010).

5. Jemison Jr., J. M. & Fox, R. H. A quick-test procedure for soil and plant tissue nitrates using test strips and a hand-held reflectometer. Commun. Soil Sci. Plant Anal. 19, 1569–1582 (1988).

6. Schmidhalter, U. Development of a quick on-farm test to determine nitrate levels in soil. J. Plant Nutr. Soil Sci. 168, 432–438 (2005).

7. Chen, K.-Y., Biswas, A., Cai, S., Huang, J. & Andrews, J. Inkjet printed potentiometric sensors for nitrate detection directly in soil enabled by a hydrophilic passivation layer. *Adv*. Mater. Technol. 9, (2024).

8. Chen, S., Chen, J., Qian, M., Liu, J. & Fang, Y. Low cost, portable voltammetric sensors for rapid detection of nitrate in soil. Electrochimica Acta 446, 142077 (2023).

9. Zhu, Y. et al. Continuous in situ soil nitrate sensors: the importance of high-resolution measurements across time and a comparison with salt extraction-based methods. Soil Sci. Soc. Am. J. 85, 677–690 (2021).

10. Li, Y., Yang, Q., Chen, M., Wang, M. & Zhang, M. An ISE-based on-site soil nitrate nitrogen detection system. Sensors 19, 4669 (2019).

11. Eldeeb, M. A., Dhamu, V. N., Paul, A., Muthukumar, S. & Prasad, S. Electrochemical soil nitrate sensor for In situ real-time monitoring. Micromachines 14, 1314 (2023).

12. Thompson, L. J. & Puntel, L. A. Transforming unmanned aerial vehicle (UAV) and multispectral sensor into a practical decision support system for precision nitrogen management in corn. Remote Sens. 12, 1597 (2020).

13. Li, Y. et al. A review of UAV remote sensing technology applications in common gramineous crops. Inf. Process. Agric. (2026) doi:10.1016/j.inpa.2026.01.009.

14. Zhang, S., Wang, X., Lin, H., Dong, Y. & Qiang, Z. A review of the application of UAV multispectral remote sensing technology in precision agriculture. Smart Agric. Technol. 12, 101406 (2025).

15. Sadoine, M., De Michele, R., Župunski, M., Grossmann, G. & Castro-Rodríguez, V. Monitoring nutrients in plants with genetically encoded sensors: achievements and perspectives. Plant Physiol. 193, 195–216 (2023).

16. DeAngelis, K. M., Ji, P., Firestone, M. K. & Lindow, S. E. Two Novel Bacterial Biosensors for Detection of Nitrate Availability in the Rhizosphere. Appl. Environ. Microbiol. 71, 8537–8547 (2005).

17. Wang, R. et al. Multiple Regulatory Elements in the Arabidopsis NIA1 Promoter Act Synergistically to Form a Nitrate Enhancer. Plant Physiol. 154, 423–432 (2010).

18. Ho, C.-H. & Frommer, W. B. Fluorescent sensors for activity and regulation of the nitrate transceptor CHL1/NRT1.1 and oligopeptide transporters. eLife 3, e01917 (2014).

19. Chen, Y.-N., Cartwright, H. N. & Ho, C.-H. In vivo visualization of nitrate dynamics using a genetically encoded fluorescent biosensor. Sci. Adv. 8, eabq4915 (2022).

20. Liu, K.-H. et al. NIN-like protein 7 transcription factor is a plant nitrate sensor. Science 377, 1419–1425 (2022).

21. Bian, C. et al. Conservation and divergence of regulatory architecture in nitrate-responsive plant gene circuits. Plant Cell 37, koaf124 (2025).

22. Johnstone, C. P. & Galloway, K. E. Supercoiling-mediated feedback rapidly couples and tunes transcription. Cell Rep. 41, (2022).

23. Yeung, E. et al. Biophysical constraints arising from compositional context in synthetic gene networks. Cell Syst. 5, 11–24.e12 (2017).

24. González-Grandío, E., Demirer, G. S., Ma, W., Brady, S. & Landry, M. P. A Ratiometric Dual Color Luciferase Reporter for Fast Characterization of Transcriptional Regulatory Elements in Plants. ACS Synth. Biol. 10, 2763–2766 (2021).

25. Witham, S. Understanding and engineering cis-regulatory functions in Arabidopsis thaliana. (University of East Anglia. School of Biological Sciences, 2023).

26. Furuhata, Y., Sakai, A., Murakami, T., Nagasaki, A. & Kato, Y. Bioluminescent imaging of Arabidopsis thaliana using an enhanced Nano-lantern luminescence reporter system. PLOS ONE 15, e0227477 (2020).

27. Li, E. et al. Vacuum and sonication treatment enables efficient transient gene expression in various monocot and eudicot plant seedlings. 2025.03.13.642903 Preprint at 10.1101/2025.03.13.642903 (2025).

28. Moreno-Giménez, E., Selma, S., Calvache, C. & Orzáez, D. GB_SynP: A Modular dCas9-Regulated Synthetic Promoter Collection for Fine-Tuned Recombinant Gene Expression in Plants. ACS Synth. Biol. 11, 3037–3048 (2022).

29. Swift, J., Alvarez, J. M., Araus, V., Gutiérrez, R. A. & Coruzzi, G. M. Nutrient dose-responsive transcriptome changes driven by michaelis–menten kinetics underlie plant growth rates. Proc. Natl. Acad. Sci. 117, 12531–12540 (2020).

30. McNickle, G. G. & Brown, J. S. When michaelis and menten met holling: towards a mechanistic theory of plant nutrient foraging behaviour. AoB PLANTS 6, plu066 (2014).

31. Ho, C.-H., Lin, S.-H., Hu, H.-C. & Tsay, Y.-F. CHL1 functions as a nitrate sensor in plants. Cell 138, 1184–1194 (2009).

32. Mus, F. et al. Genetic determinants of ammonium excretion in *nifL* mutants of azotobacter vinelandii. Appl. Environ. Microbiol. 88, e01876–21 (2022).

33. Bueno Batista, M., Brett, P., Appia-Ayme, C., Wang, Y.-P. & Dixon, R. Disrupting hierarchical control of nitrogen fixation enables carbon-dependent regulation of ammonia excretion in soil diazotrophs. PLOS Genet. 17, e1009617 (2021).

34. Bishnoi, U., Polson, S. W., Sherrier, D. J. & Bais, H. P. Draft genome sequence of a natural root isolate, bacillus subtilis UD1022, a potential plant growth-promoting biocontrol agent. Genome Announc. 3, 10.1128/genomea.696-15 (2015).

35. Powers, E. N. et al. Bidirectional promoter activity from expression cassettes can drive off-target repression of neighboring gene translation. Elife 11, e81086 (2022).

36. Bustamante, S. C. & Hartz, T. K. Nitrogen management in organic processing tomato production: nitrogen sufficiency prediction through early-season soil and plant monitoring. HortScience 50, 1055–1063 (2015).

37. Breschini, S. J. & Hartz, T. K. Presidedress soil nitrate testing reduces nitrogen fertilizer use and nitrate leaching hazard in lettuce production. HortScience 37, 1061–1064 (2002).

38. Heckman, J. In-season soil nitrate testing as a guide to nitrogen management for annual crops. HortTechnology 12, 706–710 (2002).

39. Khakhar, A. et al. Building customizable auto-luminescent luciferase-based reporters in plants. Elife 9, e52786 (2020).

40. Mitiouchkina, T. et al. Plants with genetically encoded autoluminescence. Nat. Biotechnol. 38, 944–946 (2020).

41. He, Y., Zhang, T., Sun, H., Zhan, H. & Zhao, Y. A reporter for noninvasively monitoring gene expression and plant transformation. Hortic. Res. 7, 1–6 (2020).

42. Yuan, G. et al. Expanding the application of a UV-visible reporter for transient gene expression and stable transformation in plants. Hortic. Res. 8, 234 (2021).

43. Konishi, M. & Yanagisawa, S. Roles of the transcriptional regulation mediated by the nitrate-responsive cis-element in higher plants. Biochem. Biophys. Res. Commun. 411, 708–713 (2011).

44. Maeda, Y. et al. A NIGT1-centred transcriptional cascade regulates nitrate signalling and incorporates phosphorus starvation signals in Arabidopsis. Nat. Commun. 9, 1376 (2018).

45. Carruthers, B. M., Garcia, A. K., Rivier, A. & Kacar, B. Automated laboratory growth assessment and maintenance of azotobacter vinelandii. Curr. Protoc. 1, e57 (2021).

46. Muchenje, K. T. et al. Optimized R2 retroelement complexes enable precise and efficient DNA insertion into plant genomes. 2025.08.22.671877 Preprint at 10.1101/2025.08.22.671877 (2025).

47. Hachiya, T. & Okamoto, Y. Simple Spectroscopic Determination of Nitrate, Nitrite, and Ammonium in Arabidopsis thaliana. BIO-Protoc. 7, (2017).

48. Aja-Macaya, P. et al. Nanopore full length 16S rRNA gene sequencing increases species resolution in bacterial biomarker discovery. Sci. Rep. 15, 26486 (2025).

