## Supplementary Figures for "Sentinel plants enable quantitative monitoring of bioavailable nitrate in soils and microbial environments"

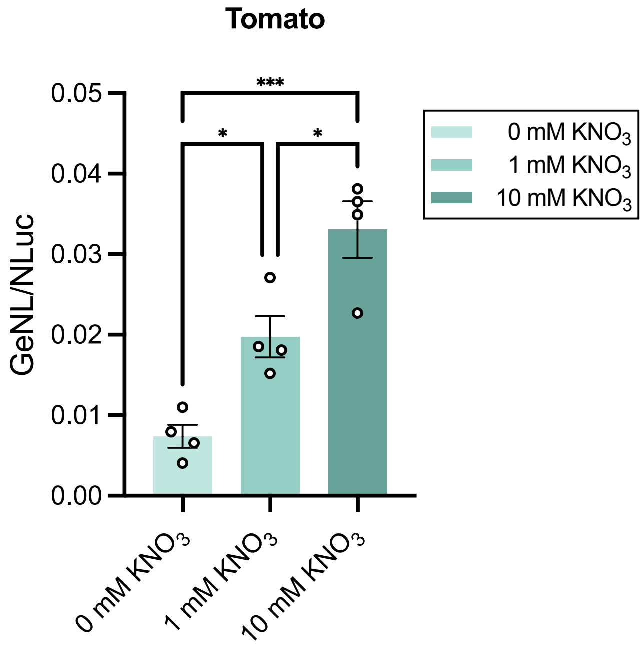


**Supplementary Figure 1.** MicroTom tomato plants transiently transformed with the NLP7 nitrate sensor plasmid respond to 0, 1, and 10 mM nitrate. Data are shown as mean ± SEM; n=4 biological replicates, each with 5 pooled seedlings. Statistical significance was determined by one-way ANOVA followed by Tukey's multiple comparisons test; *$p < 0.05$, ***$p < 0.001$.

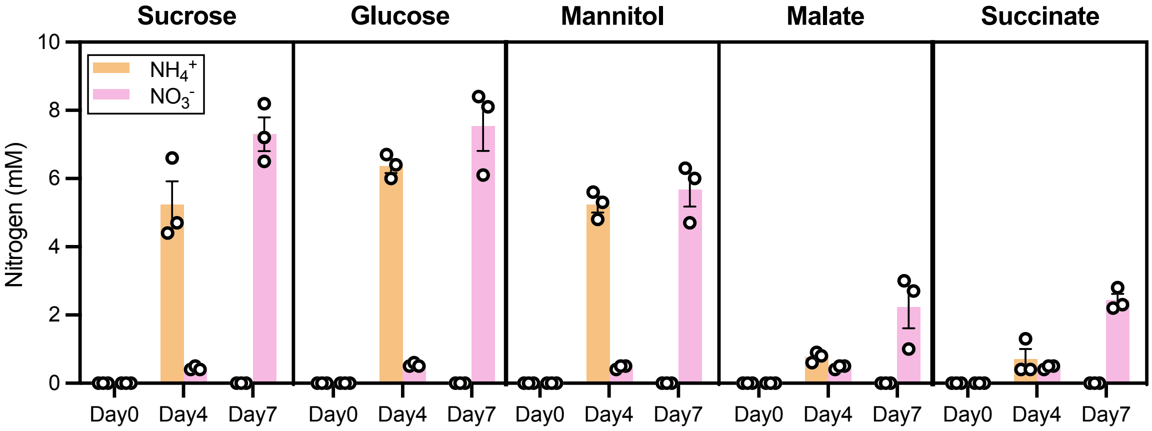


**Supplementary Figure 2.** Total ammonium and nitrate accumulation in *Azotobacter vinelandii* AV FM2–nitrifier co-cultures (nitrifiers added on day 4) grown with different carbon sources, including sugars (sucrose and glucose), the sugar alcohol mannitol, and organic acids (malate and succinate). Data are shown as mean ± SEM; n=3 biological replicates, each representing an independent culture.

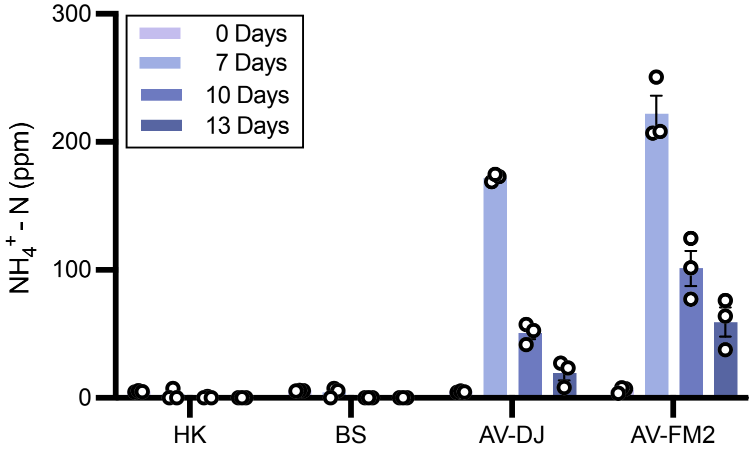


**Supplementary Figure 3.** Ammonia accumulation in peat moss inoculated with various bacterial treatments over a 13-day period. Nitrifiers (*N. europaea* and *N. winogradskyi*) were introduced to all pots at day 7. HK, heat-killed *AV-FM2* biomass control; BS, *B. subtilis* non-diazotrophic control; AV-DJ, wild-type *A. vinelandii*; AV-FM2, engineered mutant. Data represent means ± SEM; n = 3 biological replicates, each representing an independent culture.

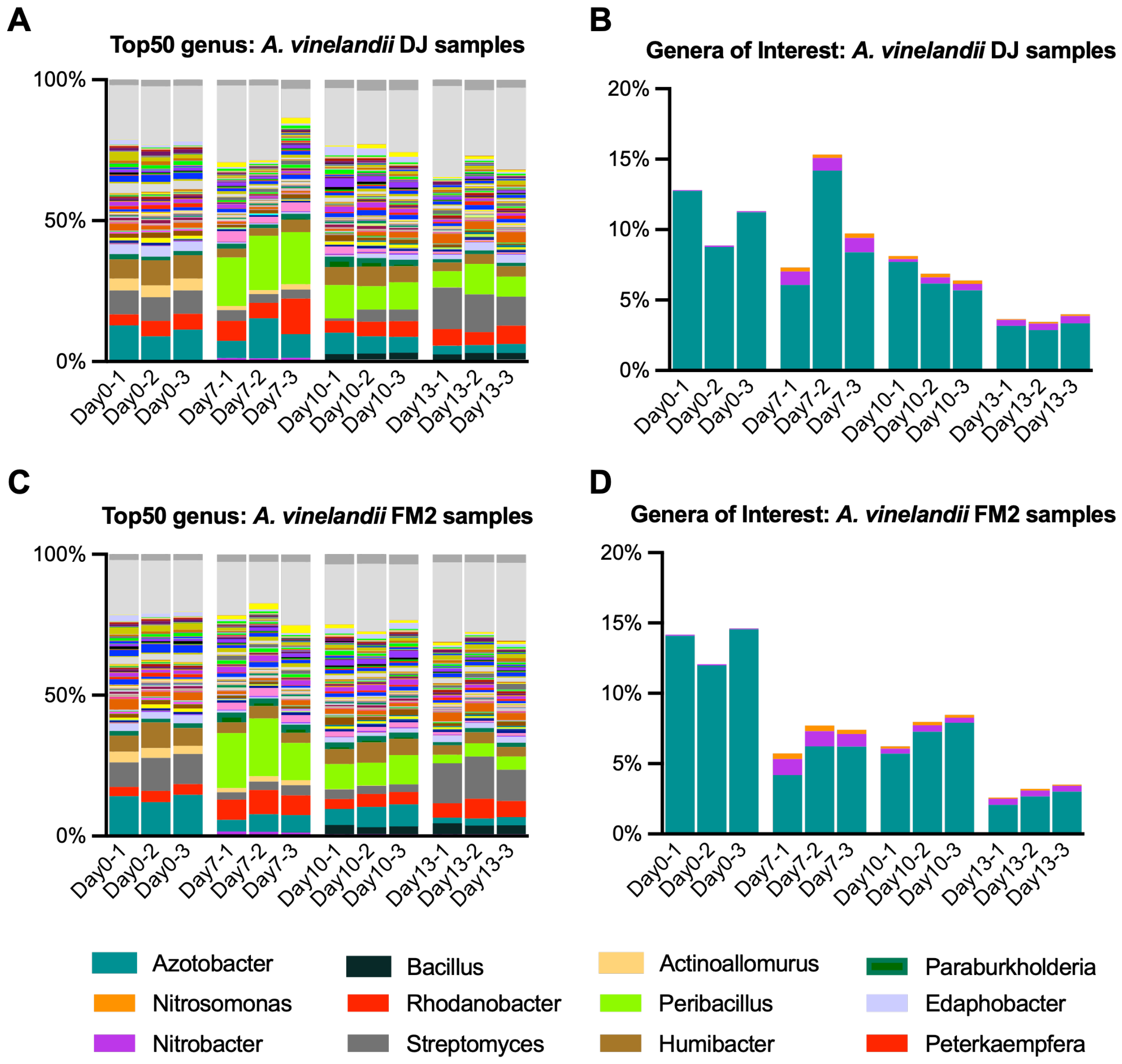


**Supplementary Figure 4. 16S rRNA gene amplicon sequencing of peat moss microcosms inoculated with synthetic nitrogen-fixing consortia.** Relative abundance of bacterial genera in AV-DJ **(A–B)** and AV-FM2 **(C–D)** treatments sampled at days 0, 7, 10, and 13. Panels A and C show the top 50 genera; panels B and D highlight selected nitrogen-cycling genera of interest. Bars represent individual biological replicates; n = 3 biological replicates, each representing an independent soil sample with different cultures.

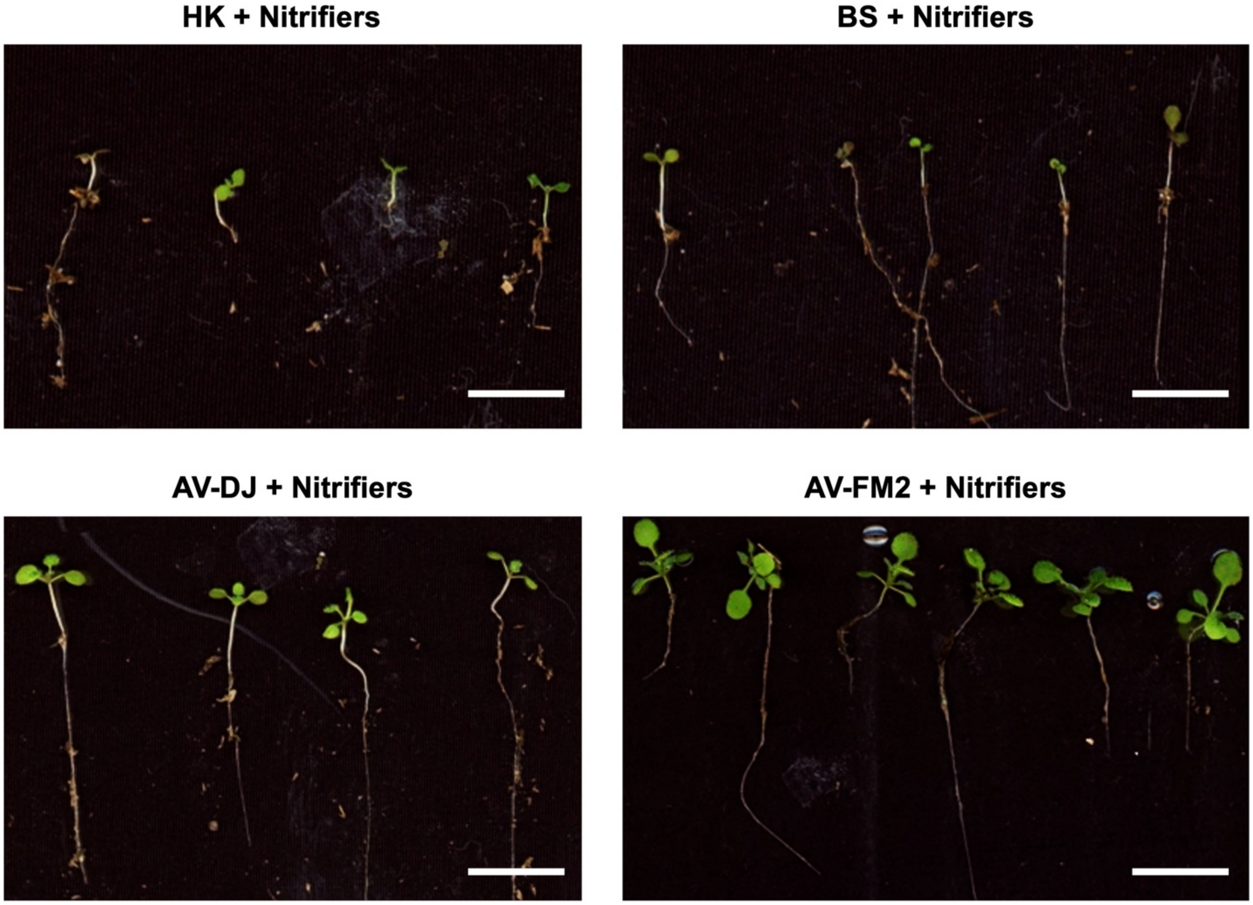


**Supplementary Figure 5. Plant biomass representative images.** Representative images of *Arabidopsis thaliana* seedlings grown for 14 days in peat moss with the indicated treatments: heat-killed (HK) consortium + nitrifiers, *B. subtilis* (BS) control + nitrifiers, *Azotobacter vinelandii* AV-DJ + nitrifiers, and *Azotobacter vinelandii* AV-FM2 + nitrifiers. 10 seedlings from each treatment were used in calculating seedling fresh weight (Fig 6C). All scale bars = 1cm.
